# Biosynthesis and chemical characterization of an intracellular red pigment of *Talaromyces islandicus* T101

**DOI:** 10.1101/2020.06.11.145821

**Authors:** Igor Vinícius Pimentel Rodrigues, Katia Regina Assunção Borges, Neurene da Cruz, Amanda Mara Teles, Marcos Antonio Custódio Neto da Silva, Rita de Nazaré Silva Alves, Marcelo Souza de Andrade, André Salim Khayat, Jaqueline Diniz Pinho, André Alvares Marques Vale, Sulayne Janayna Araújo Guimarães, Jerônimo Conceição Ruiz, Maria do Desterro Soares Brandão Nascimento, Geusa Felipa de Barros Bezerra

## Abstract

The interest in red colorants by the food industry has been increasing recently due to its wide application in many foods and beverages, and also to the carcinogenic and teratogenic effects of some synthetic dyes. Many ascomycetous fungi are able to synthesize and produce pigments, rendering them as alternative sources of natural dyes that are independent of environmental conditions. *Talaromyces islandicus* TI01 was isolated from a marine-influenced environment that has been suffering for decades from anthropogenic actions in its body of water. Broth microdilution technique was performed to analyze the antimicrobial activity. For analysis of the cytotoxic activity, the MTT [3-(4,5-dimethylthiazol-2-yl) -2,5-diphenyltetrazolium bromide] assay was conducted. The chemical analysis of the extract was performed by LC/MS (liquid chromatography coupled to mass spectrometry). The minimum bactericidal concentration (MBC) of *T. islandicus’* intracellular red pigmented extract (IRPE) for *E. coli* ATCC 25922 and *S. aureus* ATCC 25923 was 1000 μg/ml. The minimum inhibitory concentration (MIC) for *E. coli* was 250 μg/mL and for *S. aureus* 500 μg/mL, respectively, whereas for *C. tropicalis* ATCC 1369 was 62.5 μg/mL. IC50 for breast cancer cell line (MCF-7) was 45.43 ± 1.657 μg / mL. The major compounds present in the extract were: Luteoskyrin **(1)** and N-GABA-PP-V (6-[(Z)-2-Carboxyvinyl]-N-GABA-PP-V) **(2)**. The results show that IRPE from *T. islandicus* TI01 has a prominent antibacterial activity against *E. coli* and *S. aureus*, making this pigment interesting for development of new food colorants and/or conservative agents, since these bacteria are food-borne pathogens.

## Background

Filamentous fungi are known to produce a variety of secondary metabolites that are related to the resistance of these organisms to various adverse environmental factors, such as pollution, exposure to extreme temperatures, irradiation and photo-oxidation, or in ecological interactions with other organisms such as sponges, corals or other microbial communities^1^.

Pigments are among these secondary metabolites and these natural compounds are potential candidates to replace synthetic dyes, which exhibit disadvantages including toxicity to health or the environment, and mutagenic and carcinogenic properties^2^. Furthermore, many pigments have important biological properties, such as antibacterial, antifungal, antitumoral and anticholesterolemic activities, which has been leading to an increasing attention by the pharmaceutical industry^3-6^.

The *Talaromyces* genus (*Eurotiomycetes, Trichocomaceae*) was initially created to comprise the teleomorphs of biverticillate *Penicillium* species. However, according to the principle “one fungus – one name” affirmed in fungal taxonomy, which adopts a single holomorphic denomination for species presenting two different stages in their life cycle^7^, *Talaromyces* sp. now includes all species in the *Penicillium* subgenus *Biverticillium*^8^, while *Penicillium* sp. comprises the *sensu stricto* species belonging to the subgenera *Aspergilloides, Furcatum*, and *Penicillium*, their associated *Eupenicillium* teleomorphs, and species classified in a few related genera^9^.

*Talaromyces islandicus* is one of the most destructive and harmful fungi that affect rice in storage, causing the yellowing of rice^10-11^ and is also able to produce mycotoxins such as cyclochlorotine, islanditoxin, erythroskyrine and luteosyrin, which are hepatotoxic and carcinogenic agents^12^. Luteoskyrin is in the IARC Group 3 carcinogen and yield of this substance is the highest among the metabolites of *T. islandicus*^13^.

Due to the increasing resistance acquired by microrganisms to antibiotic and antifungal drugs commonly used in the clinical setting, antimicrobial agents of natural origin are being used to treat microbial infections, thus saving millions of lives^14^. Moreover, cancer is a major cause of morbidity and mortality in developing and developed countries, and it is well documented the resistance of many cancerous cells to antitumoral drugs, which consists in an obstacle for cancer eradication, mainly due to a failure of the damaged cells to activate apoptotic pathways. It is also reported a protective effect of autophagy of cancer cells, making it harder to develop an effective strategy to counteract cancer cell growth and improving the response to therapy^15^.

In an attempt to discover new molecules with antimicrobial and antitumoral activities, we investigated, in the present study, the antifungal, antibacterial and cytotoxic activities of an intracellular red pigment extract (IRPE) produced by *Talaromyces islandicus* TI01 isolated from a soil influenced by a marine environment (State Park of Jansen’s Lagoon located in São Luís, Maranhão, Brazil) that has been suffering for decades from the anthropogenic actions of the nearby houses and commercial buildings, against *Candida albicans* ATCC 10231, *Candida tropicalis* ATCC 1369, *Escherichia coli* ATCC 25922 and *Staphylococcus aureus* ATCC 25923 as well as its cytotoxic activity against breast cancer cell line (MCF-7). We also tried to characterize the chemical compounds present in *T. islandicus’* TI01 IRPE, comparing them to substances already described in the scientific literature.

## Results

### Yield of *T. islandicus’* TI01 IRPE

From a total of 1.5 liters of Potato-Dextrose Broth where the fungus was cultivated, it was obtained a total of 6.660 grams of fungal biomass and 0.857 grams of intracellular pigment, which was contained in the mycelium of the fungus (Figs 1 and 2)

**Figure 1.**
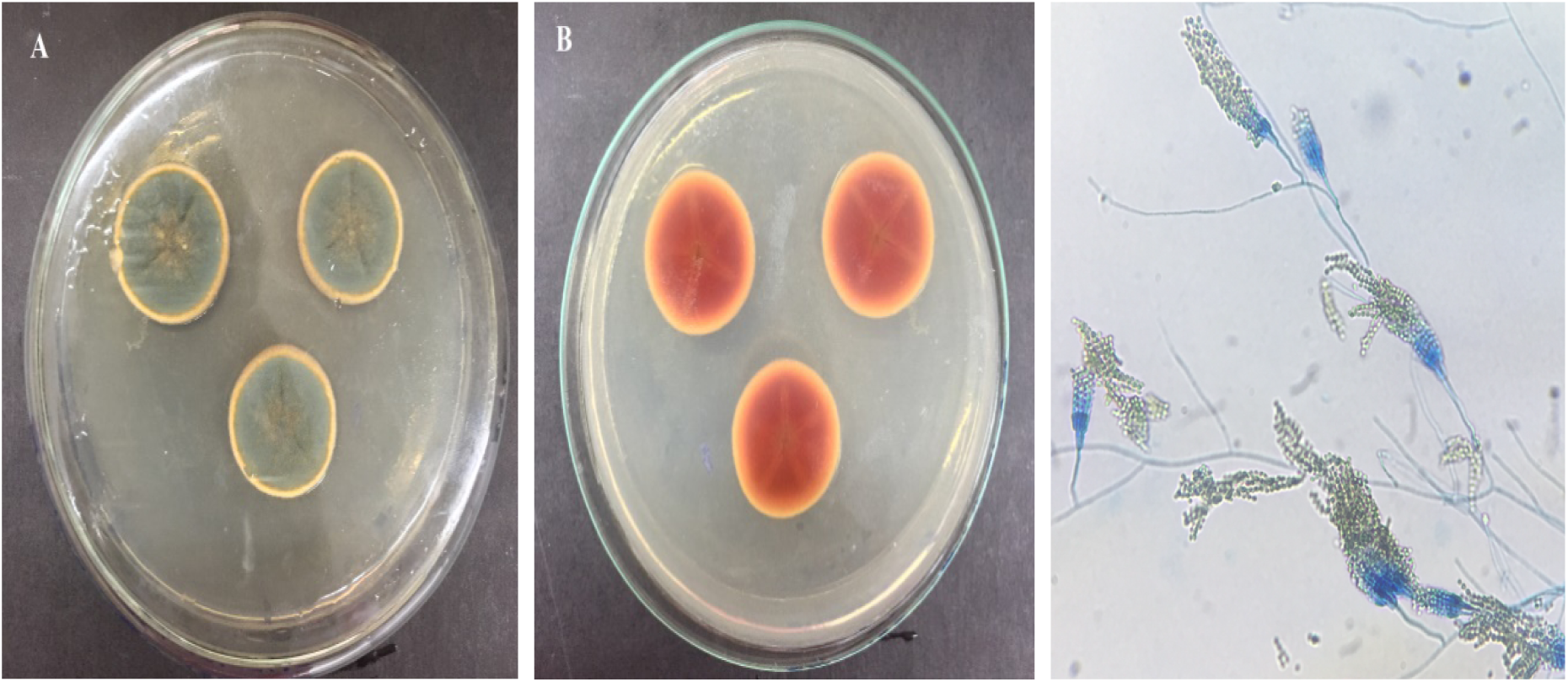
Macroscopy and microscopy of *Talaromyces islandicus*. 1A: Obverse of *Talaromyces islandicus* colonies isolated on Sabouraud-Dextrose Agar. 1B: Reverse of *T. islandicus* colonies isolated on Sabouraud-Dextrose Agar. 1C: Microscopy of T. *islandicus*

**Figure 2.**
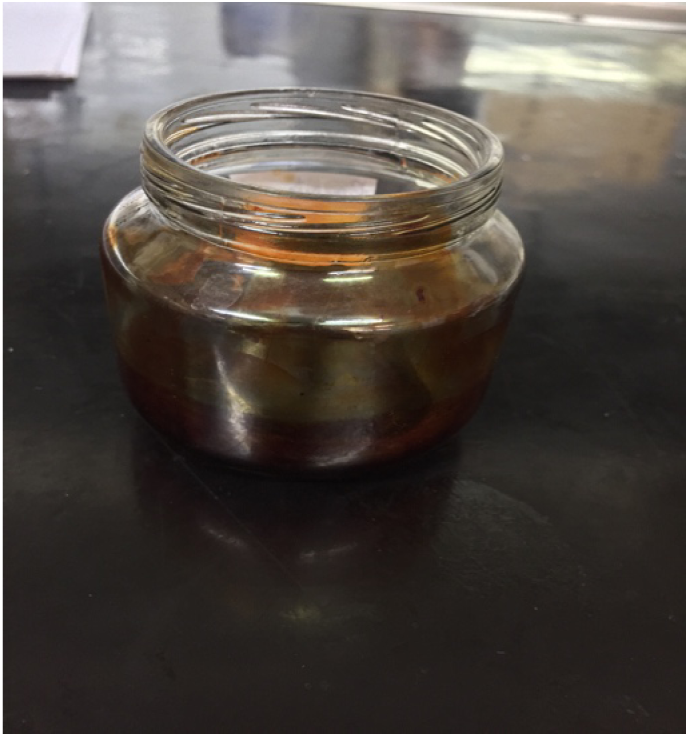
Red pigment obtatined from T. islandicus

### Chemical characterization of *T. islandicus*’ TI01 IRPE

Figure 3 shows the chemical profile of the extract from a chromatographic run with a duration of 68 minutes at a wavelength of 254 nm, obtaining eight peaks where most of the compounds present in the extract were concentrated. The m/z, retention times and area in percentage (%) are illustrated in table 1. The spectra of each compound corresponding to the obtained peaks are illustrated in the Supplementary Material.

**Table 1.** Probable compounds present in *T. islandicus*’ IRPE.

| Peak | m/z | RT | Probable Compounds | Area % | Chemical classes | References |
| --- | --- | --- | --- | --- | --- | --- |
| 1 | 481,19 | 8.4 | HHDP-glucose | 0,185709 | Hydrolyzable Tannin | Robbins et al<sup>16</sup> |
| 2 | 302,97 | 12.3 | Ganoderic acid C2 | 0,326711 | Triterpenoid | Cao et al<sup>17</sup> |
| 3 | 649,04 | 20.1 | Galloyl-HHDP-gluconate (lagerstannin C) isomer | 2,992329 | Tannin | Mena et al<sup>18</sup> |
| 4 | 589,09 | 45.5 | Rhamnosyl-hexosyl-acyl-quercetin | 0,660952 | Flavonoid | Said et al<sup>19</sup> |
| 5 | 573,05 | 50.3 | Luteoskyrin | 74,84014 | Naphtalene | Peng et al<sup>12</sup> |
| 6 | 595,01 | 59.2 | Pelargonidin-3,5-diglucoside | 1,458194 | Anthocyanidin | Barnes; Shug<sup>20</sup> |
| 7 | 931,46 | 60.1 | Notoginsenoside-R1 | 0,86856 | Ginsenoside | Sun et al<sup>21</sup> |
| 8 | 454,20 | 61.5 | N-GABA-PP-V (6-[(Z)-2-Carboxyvinyl]-N-GABA-PP-V) | 18,66741 | Azaphilone polyketide | Venkatachalam et al<sup>22</sup> |

**Figure 3.**
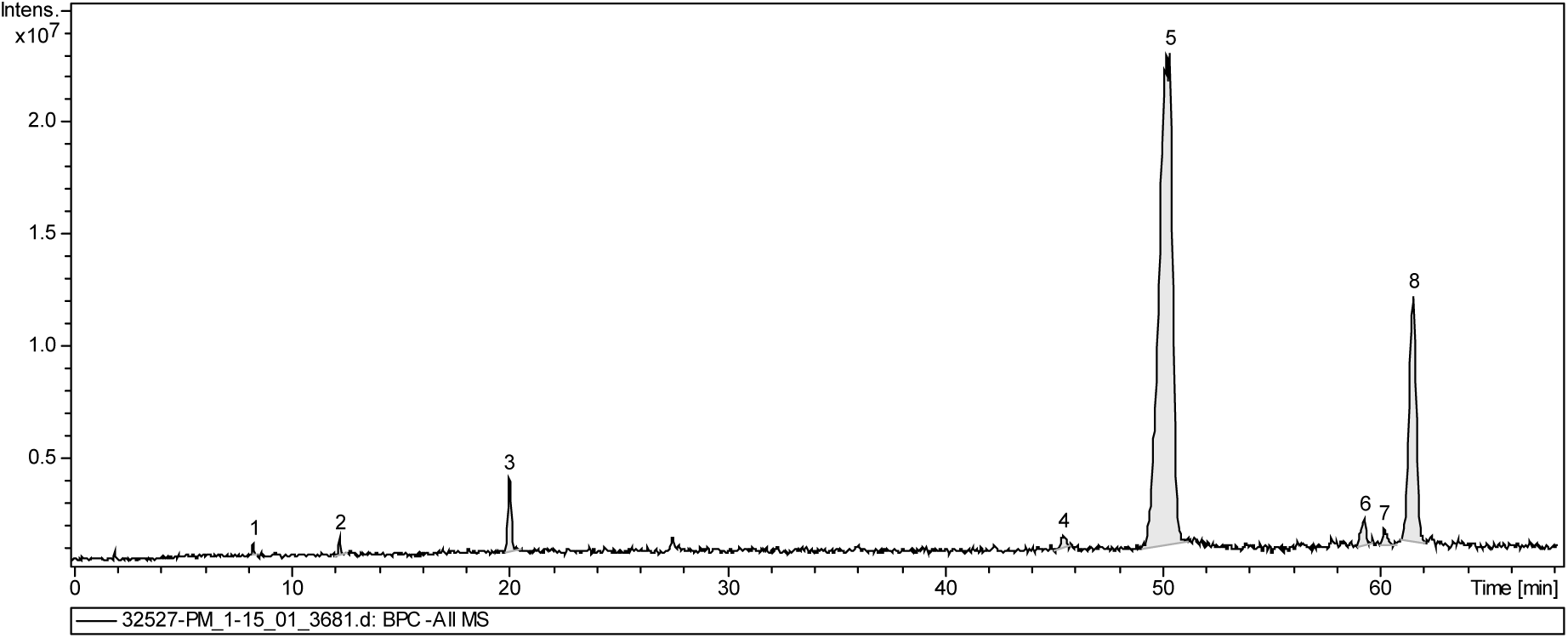
Chromatographic profile of *T. islandicus’* IRPE (wavelength of 254 nm).

Peak 1 was detected at 8.4 minutes, peak 2 at 12.3 minutes, peak 3 at 20.1 minutes, peak 4 at 45.5 minutes, and peak 5 at 50.3 minutes, peak 6 detected at 59.2 minutes, peak 7 to 60.1 minutes, peak 8 to 61.5 minutes.

### Molecular identification of the fungal strain

The size of the small subunit ribosomal RNA gene, internal transcribed spacer 1 (ITS 1) and 5.8 S ribosomal RNA gene region amplified in this work was 328 base pairs (bp) (Figure 4). The nucleotide BLAST search result showed that the amplified sequence was most similar (98.43%) to the sequence of *Talaromyces islandicus* strain CBS 178.68 and the e-value was of 3e^-161^. The strain is deposited on the Collection of Fungi of the Laboratory of Mycology, Nucleus for Basic and Applied Immunology, Center for Biological and Health Sciences, Federal University of Maranhão.

**Figure 4.**
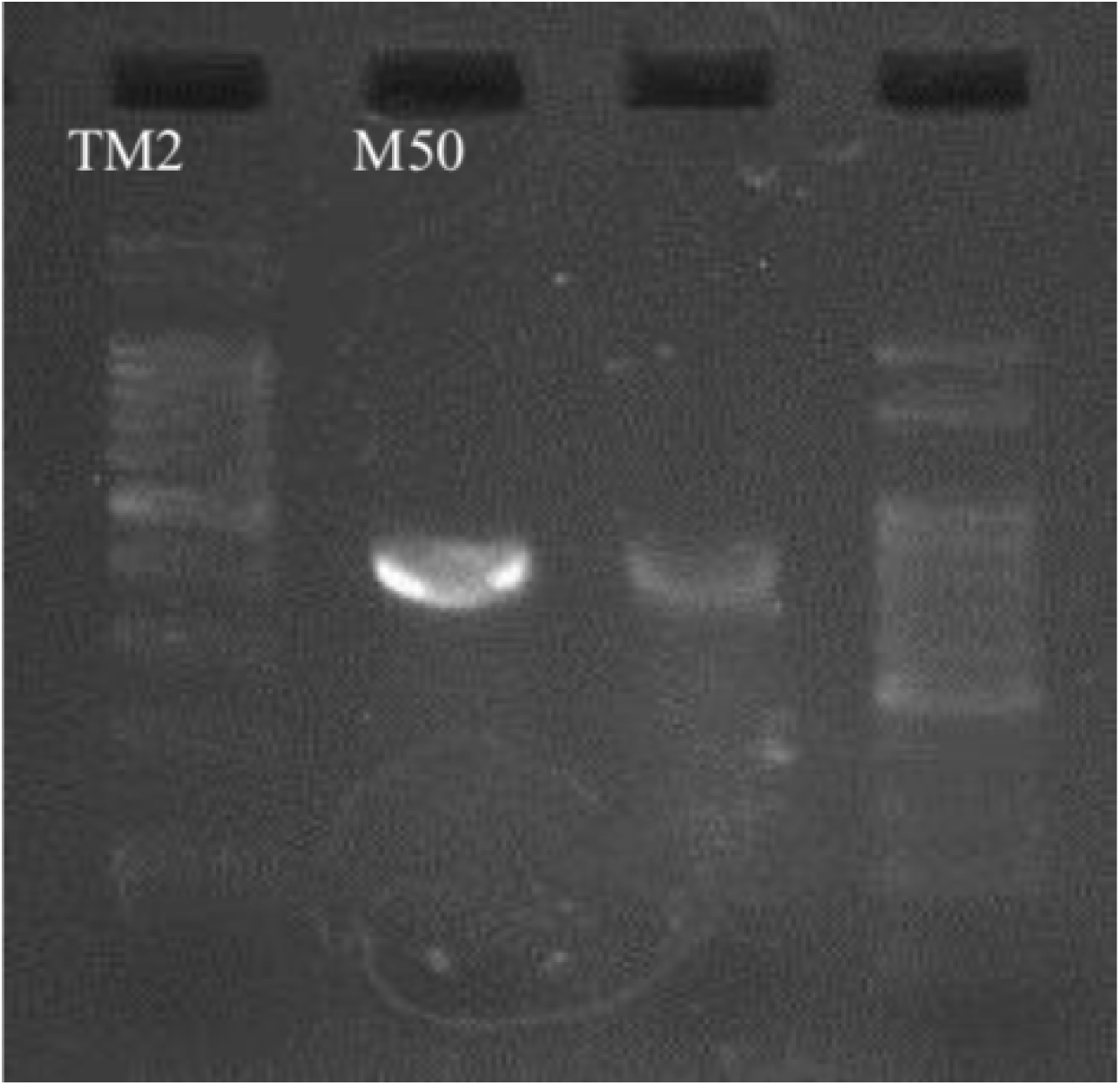
PCR amplification of DNA from *T. marneffei* with specific primers TM1 and TM2. M100: Size marker of 100-bp-ladder standard DNA. M50: size marker of 50-bp-ladder standard DNA

### Biological activities of *T. islandicus*’ IRPE

#### *T. islandicus* is effective against *E. coli*

The intracellular red pigmented extract (IRPE) from *T. islandicus* TI01 was bactericidal against *E. coli* ATCC 25922 and *S. aureus* ATCC 25923 just at 1.000 μg/mL. MBC against both bacteria was 3x the MIC. MIC (Minimum Inhibitory Concentration) against *E. coli* was 250 µg/mL (p = 0.0005) and 500 µg/mL against *S. aureus* (p = 0.0078) since the log of colony forming units (CFUs) showed a significant statistical difference compared to the negative control (Figure 5). There was no growth of colonies in the positive controls (4 μg/mL gentamycin for *E. coli)* and 8 μg/mL vancomycin for *S. aureus*).

**Figure 5.**
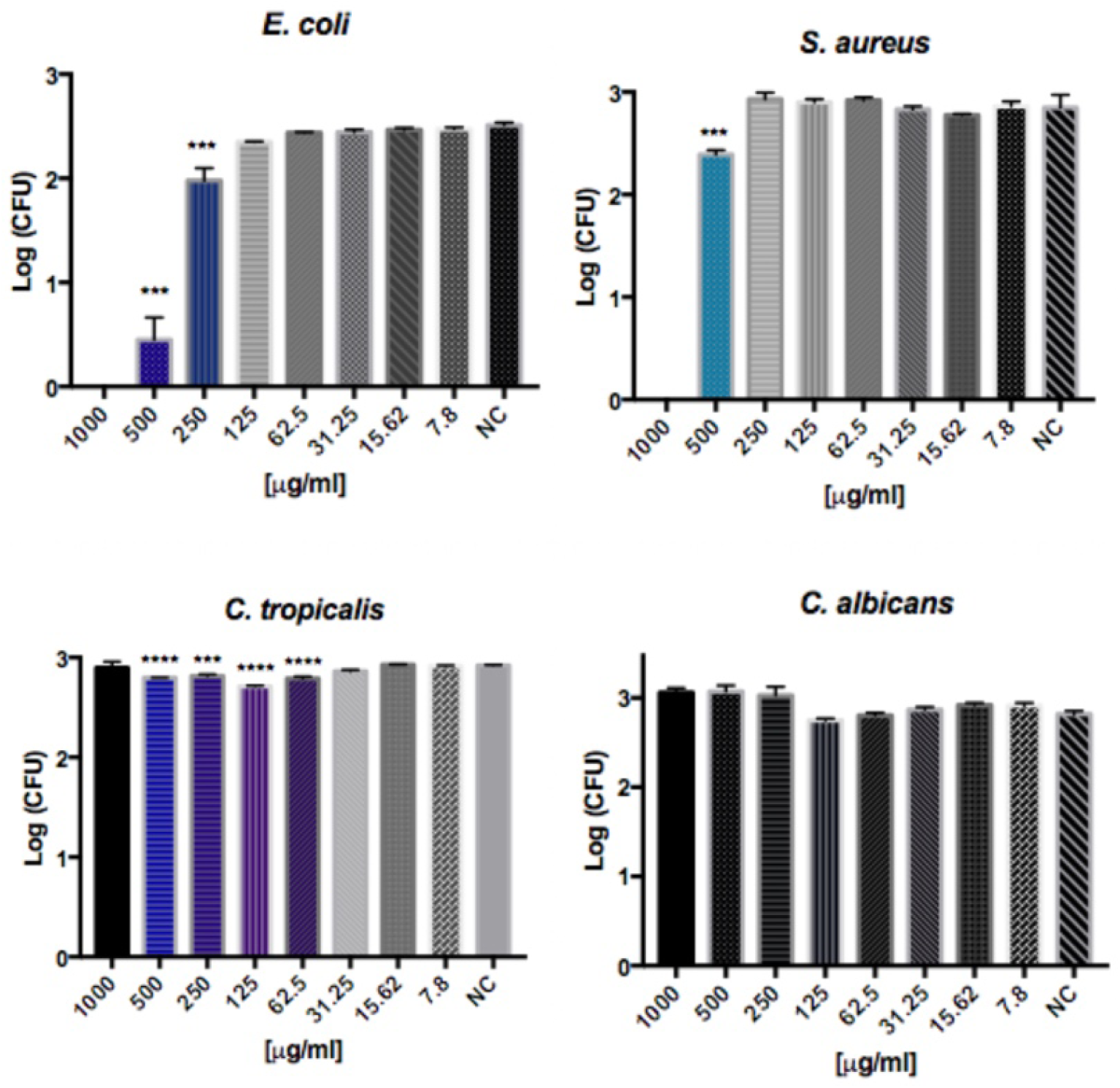
Logarithm of the number of Colony Forming Units (CFUs) of the tested microorganisms in the presence of decreasing concentrations of IRPE from *T. islandicus* TI01. From the top left to the bottom right: *E. coli* ATCC 25922; *S. aureus* ATCC 25923; *C. tropicalis* ATCC 1369 and *C. albicans* ATCC 10231. Error bars represent standard deviation. Statistical p values (represented by **\*\***\* or \*\***\***\*) indicates concentrations that are significantly different from control. **\*\***\* p < 0.05; \*\*\***\*** p < 0.001.

MIC value for *C. tropicalis* ATCC 1369 was 62.5 μg/mL (p < 0.0001) (Figure 5). It is noteworthy that at the concentration of 125 μg/mL, the log of CFUs (2.71) was lower compared to the other concentrations that also showed a significant statistical difference in relation to the negative control (500-62.5 μg/mL), showing that this concentration presented a greater fungistatic activity compared to the others. *T. islandicus*’ IRPE was not active against *C. albicans* ATCC 10231. There was no growth of colonies in the positive controls (AMB at 16 µg/mL).

#### *T. islandicus* red pigmented extract induces breast cancer cell death

The intracellular red pigmented extract (IRPE) from *T. islandicus* TI01 promoted breast cancer cell’s viability reduction after 72 hours of treatment at the lowest concentration tested: 0.25 µg/mL, compared to the other treatments, with a mean cellular viability of 63.13%, although it showed no statistical difference compared to the negative control (p=0.0713) (Figure 6). The mean IC50 value was 45.43 **±** 1.657 µg/mL.

**Figure 6.**
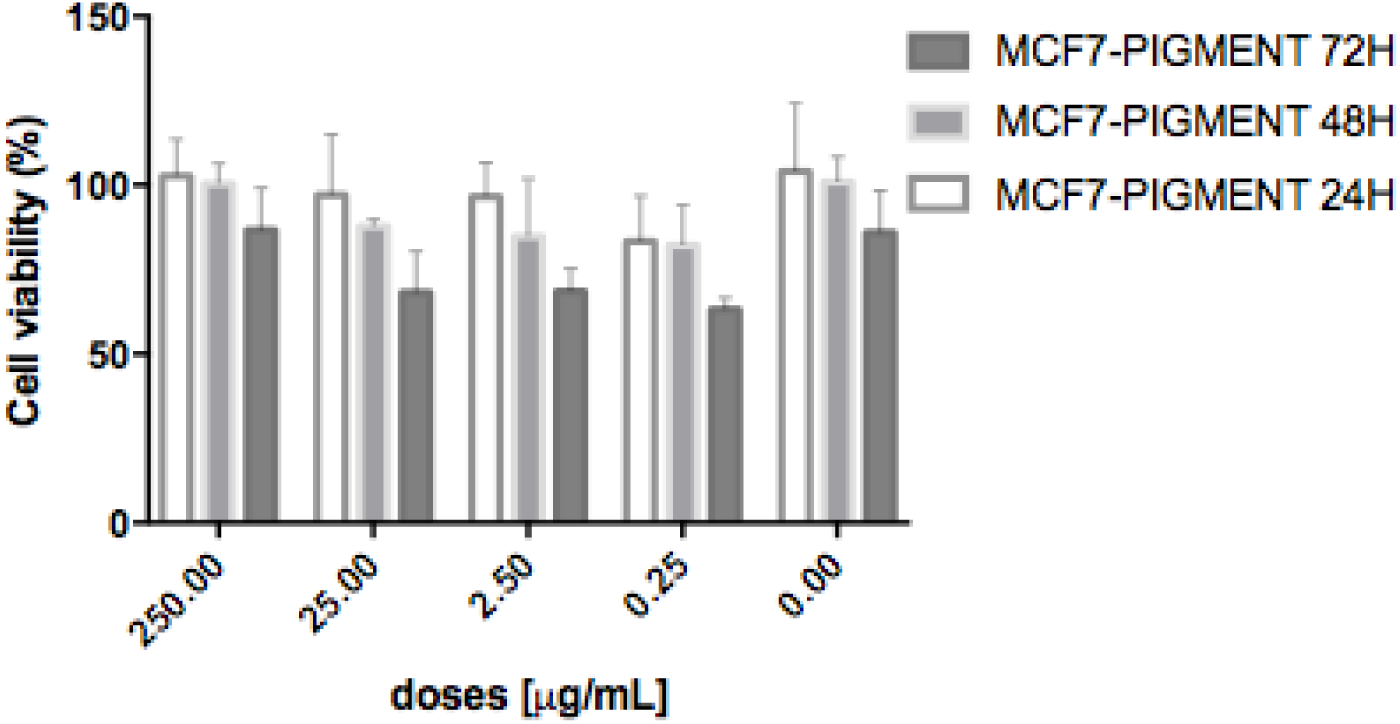
Reduction of MCF-7 cells’ viability in all treatments, with a greater cell viability reduction after 72 hours of treatment at the dose of 0.25 μg/mL. 0.00: negative control.

### *T. islandicus* pigmented extract has antioxidant activity

Antioxidant activity (%) increased proportionally with extract concentration,, producing an EC_50_ (concentration required to achieve 50% antioxidant activity) of 84,3201µg/mL (Figure 7).

**Figure 7.**
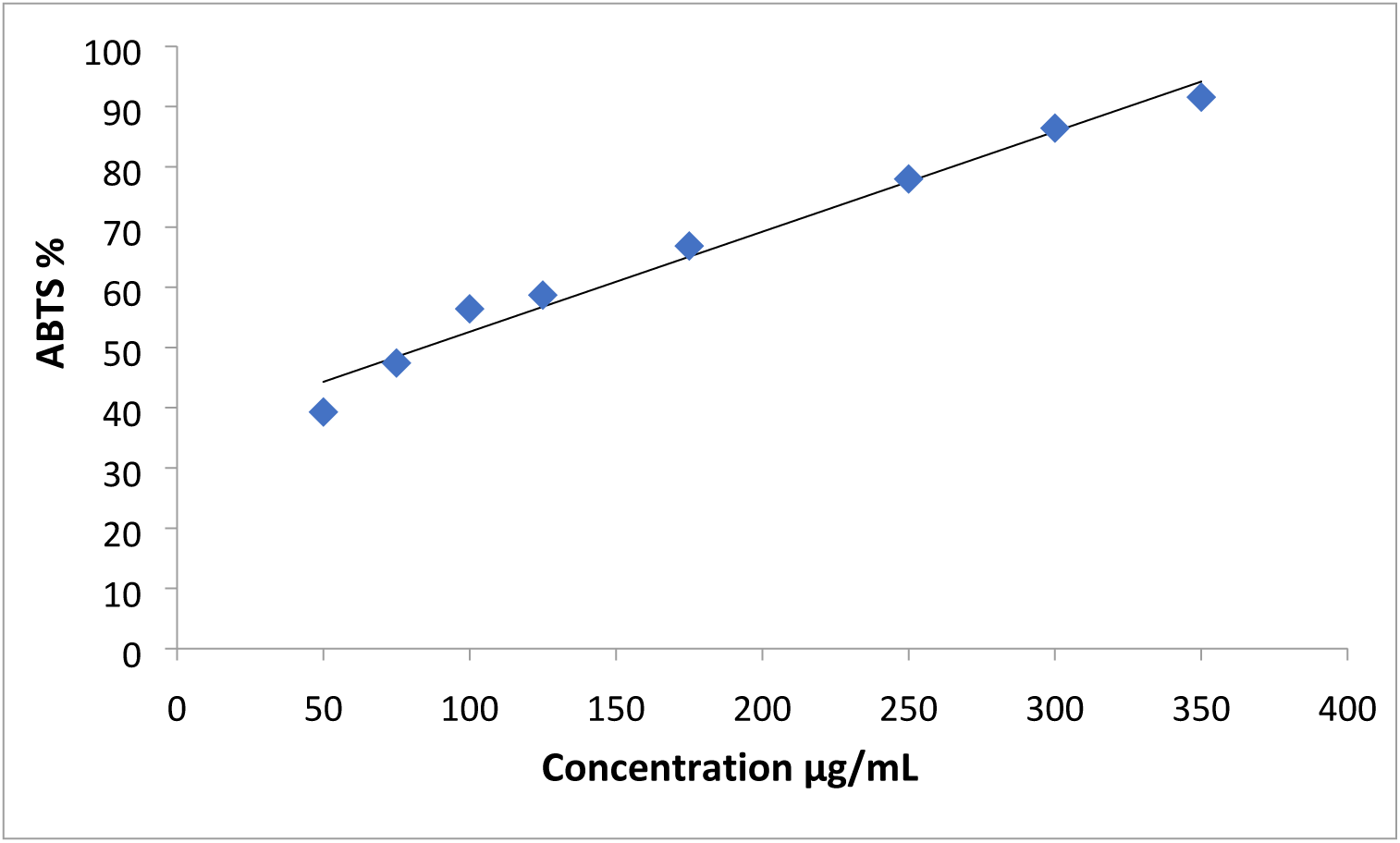
Antioxidant activity of the red pigmented extract of *T. islandicus* by ABTS method.

## Discussion

In this study, we used ethyl acetate as single solvent to extract *T. islandicus*’ TI01 red pigmented extract. According to Venkatachalam et al^22^, this is the best solvent to be used to extract pigments from fungi belonging to the *Talaromyces* genus followed by ethanol because it is able to extract the major pigmented compounds from these fungi.

Yield of luteoskyrin is the highest among metabolites of *T. islandicus*, which corroborates the finding of this study, where we tentatively identified that luteoskyrin was the majoritarian compound, with an area of 74.84%. In addition, scientific literature reports that this substance inhibits replication, transcription, and DNA repair in bacteria, yeast, and animal cells and forms chelates with nucleic acids^23^, which may be responsible for the bactericidal acitivity of *T. islandicus’* TI01 IRPE at the highest concentration tested.

*Talaromyces* species are known to produce intra and extracelullar yellow, orange and red pigments. Azaphilone polyketide pigments like mitorubrins (mitorubrin, mitorubrinol, mitorubrinol acetate and mitorubrinic acid) and the *Monascus* red pigments (N-glutaryl monascorubramin, N-glutarylrubropunctamin, monascorubramine, monascin, PP-R and others)^24^ are responsible for the red pigments and are produced in different ratios and amounts between different isolates and species. N-GABA-PP-V (6-[(Z)-2-Carboxyvinyl]-N-GABA-PP-V) is an azaphilone polyketide and it was tentatively identified as the second majoritarian compound present in *T. islandicus’* TI01 IRPE, which corroborates these data found in the literature.

Furthermore, literature affirms that phenolic compounds and flavonoids possess anti-bacterial activity^25,26,27,28^. Galloyl-HHDP-gluconate (lagerstannin C) isomer is a phenolic compound and it was tentatively identified as the third majoritarian compound present in *T. islandicus*’ TI01 IRPE. Rhamnosyl-hexosyl-acyl-quercetin is a flavonoid, and it it was tentatively identified in the present extract. It is also known that the presence of gallic or galloyl moiteties to chemical compounds induces damage to the bacterial membrane^29^, while flavonoids inhibit nucleic acid synthesis, cytoplasmic membrane function, energy metabolism, attachment and biofilm formation, porin on the cell membrane, alters the membrane permeability and attenuates the pathogenicity^30^. Therefore, these substances may have also contributed to the relevant anti-bacterial activity exhibited by the present extract.

Soumya et al^31^ found an IC50 value of a solid red pigment of oily nature produced by *Fusarium chlamydosporum*, also a filamentous fungus, of 62 μg/mL against MCF-7. The tested substance in this work showed a lower IC50 value (45.43 μg/mL) against the same tumor cell line, revealing its cytostatic activity in MCF-7 cels.

The intracellular red pigment extract (IRPE) from *T. islandicus* TI01 showed bactericidal activity against two human pathogenic bacteria: *S. aureus* and *E. coli* at the highest concentration tested. Furthermore, IRPE exhibited a weak cytotoxic activity on MCF-7 cell line. In view of the results found, future *in vivo* studies should be performed to verify if *T. islandicus*’ IRPE does not exert any cytotoxic effect on an animal model. It is important to highlight that these pathogens are related to food contamination making this pigment a potential candidate to be used as a food dye and conservative agent.

## Material and Methods

### Collection site

*T. islandicus* TI01 was isolated from the soil of a polluted environment in São Luís, Maranhão, Brazil, named Jansen Lagoon State Park, whose geographic coordinates are: 2º 30’ 13” S 44º 17’ 53” W.

The samples were obtained at a depth of up to 20 cm, with the aid of a sterile spoon according to Silva et al^32^ with modifications. Then, they were placed in a zipped plastic bag and transported to the Laboratory of Mycology (Nucleus for Basic and Applied Immunology/Department of Pathology/Center of Biological and Health Sciences/ Federal University of Maranhão), where they were processed. The fungal strain is deposited in the Collection of Fungi of the Federal University of Maranhão.

### Fungus identification

The isolate was identified by analysis of its morphological characteristics and ITS (internal transcribed spacer) gene sequence, which has been submitted to GenBank (accession number: MN831880). DNA was extracted using DNEasy Plant Mini Kit from Qiagen. The pair of primers designed was: 5’ CGT AAC AAG GTT TCC GTA GGT 3’ (forward) and 5’GTG CTT GAG GGC AGA AAT GA 3’ (reverse). A BLAST search result indicated that the sequence is almost the same (98.48%) to the sequence of *Talaromyces islandicus* strain CBS 178.68.

The pair of primers cited above was subjected to automatic sequencing using the ABI PRISMTM 310 Big Dye Terminator v3.1 Matrix Standards kit (Applied Biosystems). With the help from a Phred/Phrap/Consed pack^333,34,35,36^ it was generated a consensus sequence that was used for the molecular identification of the fungal strain.

### Natural Product Extraction

The fermentation was carried out statically, in the dark, in Potato-Dextrose Broth (PDB) (Kasvi, pH 5.1 ± 0.2 at 25° C) in 1,000 mL Erlenmeyers flasks containing 500 mL of the liquid medium in triplicate. The flasks were incubated for 21 days at 25° C. Thereafter, 300 ml of Ethyl Acetate was added to 500 ml of the fermented broth and left overnight for decantation. The mycelium was then separated from the liquid medium by filtration on Whatman paper no. 4. Then, 300 ml of Ethyl Acetate was added to the fungal biomass to provide the intracellular pigment. The solutions were placed in a separation funnel to obtain purer solutions. They were then concentrated in a rotaevaporator and stored at 4°C for further analysis of the biological activities *in vitro*.

### Chemical characterization of *T. islandicus*’ TI01 IRPE

After solubilization, the extract was analyzed by high performance liquid chromatography (HPLC) using a Shimadzu® chromatograph (ShimadzuCorp. Kyoto, Japan), consisting of a solvent injection module with a LC-20ADShimadzu detector UV-Vis pump (SPDA-20A) - Shimadzu. The column used was SupelcoAscentisC-18 (250×4.6mm - 5um). The elution solvents used were A (water) and B (methanol).

The established elution gradient started from 5% B for 1 minute, 30% B for 14 minutes, 60% B for 15 minutes, 60% B for 30 minutes, 70% B for 30 min, 70 % B for 7 minutes, 5% B for 1 minute rebalancing the column, using a flow rate of 1.0 mL / min, using oven temperature of 40 ° C. The sample injection volume was 20 μL. The data were collected and processed using the LCSolution software (Shimadzu). Under the conditions employed, a baseline separation was obtained for the main components of the sample in a chromatographic run of 68 minutes with a wavelength of 254nm.

### Antimicrobial Assay

Antimicrobial activity evaluation against four human pathogens (*E. coli, S. aureus, C. albicans*, and *C. tropicalis*) was carried out by microplate assay in triplicate, according to the National Comitee On Clinical Laboratory Standards^37,38^. To determine MBCs and MICs against these microrganisms, each well containing the pigment extract was plated on Petri dishes and the log of Colony Forming Units was calculated and compared to the negative and positive controls. Gentamycin at a concentration of 10 μg/ml was used for *E. coli* ATCC 25922 and vancomycin at 8 μg/ml was used for *S. aureus* ATCC 25923. Amphotericin at 16 μg/ml was used for both yeasts (*C. albicans* ATCC 10231 and *C. tropicalis* ATCC 1369), as positive controls. Negative controls consisted of *T. islandicus*’ TI01 IRPE added to the culture medium (Mueller-Hinton Broth for bacteria and BHI Broth for yeasts) at concentrations that ranged from 1.000 to 7.8 μg/ml and microorganisms’ inocula standardized at 0.5 MacFarland scale.

### Cell viability assay

Antitumoral activity was evaluated by the MTT assay [3-(4,5-dimethylthiazol-2-yl) −2,5-diphenyltetrazolium bromide]. Briefly, MCF-7 cells (2.0 × 10^4^ per well plus DMEM medium) were seeded onto a sterile 96-well plate. After a 24 hr incubation, the medium was replaced with 100 μL of fresh FBS-free medium which contained IRPE at concentrations of 250, 25, 2.5 and 0.25 μg/mL. Negative control, containing only medium with 10% SFB and tumoral cells. The plate was incubated for 24, 48 and 72 h at 37 C in 5% CO_2_ and then the media were discarded. Afterwards, the cells were stained with 100 μL of MTT solution (0.5 mg/ml) at 37° C for 4 h^39^. Thereafter, the supernatant was aspirated, and 100 μL of ethyl alcohol was added to dissolve the formazan. The optical density at 540 nm (OD540) was determined. The experiment was performed in triplicate.

The viability of the cells was quantified in percentage, using the following equation:

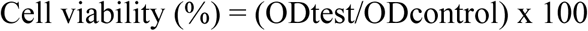

where ODcontrol is OD570 in the negative control wells (cells incubated with media only) and ODtest is the OD570 of the cells exposed to IRPE.

### Antioxidant activity by 2,2’-azinobis-[3-ethylbenzthiazoline-6-sulfonic acid] decolorization assay (ABTS)

The antioxidant activity of the extracts was evaluated using 1,2’-azinobis-[3-ethylbenzthiazoline-6-sulfonic acid (ABTS), according to methods described by Re et al^40^. For a range of extract concentrations (50 a 350 µg/mL), reaction mixtures with ABTS were prepared.

The ABTS • + radical was prepared by reacting 5.0 mL of a 3840 µg / mL ABTS solution with 88 µL of the 37,840 µg / mL of potassium persulfate solution. The mixture was left in a dark environment for 16 hours. After radical formation, the mixture was diluted in ethanol (approximately 1:30 v/v) until an absorbance of 0.7 to 734 nm was obtained.

From the concentrations of the extracts (50 to 350 µg / mL) the reaction mixture was prepared with the cation radical ABTS. In a dark environment, an aliquot of 30 µL of each concentration of the extracts was transferred in test tubes containing 3.0 mL of the radical ABTS cation and homogenized in a tube shaker. After 6 minutes, the absorbance of the reaction mixture was read on a spectrophotometer. in length of 734 nm. The analyzes were performed in triplicate and the capture of the free radical was expressed as a percentage of inhibition (% I) of the ABTS radical cation according to Babili et al^41^

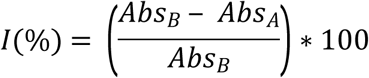

Where: I%: Percentage of inhibition of the ABTS radical

AbsA: absorbance of the sample (extract) after 6 min

AbsB: absorbance at 734 nm of the ABTS radical solution

From the data obtained, the efficient concentration or EC50% was calculated, defined as the concentration of the sample required to sequester 50% of the ABTS radicals. The extract is considered active when it has an EC50% <500 µg / mL^42^

### Statistical analysis

Results were analyzed using one-way analysis of variance (ANOVA) and applying Dunnett’s multiple comparisons post-test, using GraphPad Prism 6 (Graphpad Software, CA, USA).

## Data Availability

The data used to support the findings of this study are available from the corresponding author upon request.

## Acknowledgments

The authors would like to express their gratitude to the Federal University of Maranhão (UFMA) for allowing the use of the Applied and Basic Immunology Center (NIBA) facilities, and to the Foundation for Research and Scientific and Technological Development of Maranhão - FAPEMA that helped in financing and concretization of this study.

The authors would also like to express their gratitude to Ana Paula. S. A. dos Santos, for allowing the use of Laboratory of Immunology Applied to Cancer (LIAC) facilities of the Federal University of Maranhão (UFMA).

## Funding Statement

This work was supported by the Foundation for the Support of Research and Scientific and Technological Development of Maranhão – FAPEMA [grant number 31/2016] that financed the materials needed for the execution of this research.

## Author Contributions

I.V.P.R. prepared the manuscript and performed the experiments for the extraction and antimicrobial evaluation; K.R.A.B and R.N.S.A helped in the antimicrobial evaluation; N.C. helped in the extraction of the red pigmented extract from *T. islandicus* TI01; A.M.T. helped in the chemical analysis; M.S.A., J.D.P. JCR and A.S.K. helped in the molecular identification of the fungal strain; A.M.V, MACNS and S.J.A.G helped in the antitumoral evaluation and M.D.S.B.N. and G.F.B.B. supervised the research work and revised the manuscript.

## Conflict of interest

All authors revised the manuscript and there is no conflict of interest.

